# Computational Fluid Dynamics Modeling of Cough Transport in an Aircraft Cabin

**DOI:** 10.1101/2021.02.15.431324

**Authors:** Malia Zee, Angela C. Davis, Andrew D. Clark, Tateh Wu, Stephen P. Jones, Lindsay L. Waite, Joshua J. Cummins, Nels A. Olson

**Affiliations:** The Boeing Company, 100 N Riverside, Chicago, IL 60606

## Abstract

To characterize the transport of respiratory pathogens during commercial air travel, Computational Fluid Dynamics simulations were performed to track particles expelled by coughing by a passenger assigned to different seats on a Boeing 737 aircraft. Simulation data were post-processed to calculate the amounts of particles inhaled by nearby passengers. Different airflow rates were used, as well as different initial conditions to account for random fluctuations of the flow field. Overall, 80% of the particles were removed from the cabin in 1.3 to 2.6 minutes, depending on conditions, and 95% of the particles were removed in 2.4 to 4.6 minutes. Reducing airflow increased particle dispersion throughout the cabin but did not increase the highest exposure of nearby passengers. The highest exposure was 0.3% of the nonvolatile mass expelled by the cough, and the median exposure for seats within 3 feet of the cough discharge was 0.1%, which was in line with recent experimental testing.

## Introduction

The COVID-19 pandemic has created a global health and economic crisis. While vaccines reported to have high short-term efficiencies against the early variants of SARS-CoV-2 have been approved by regulatory authorities across the world and are being distributed widely in many countries, the ability to reach herd-immunity levels of vaccination is beginning to be considered unlikely, and uncertainty remains about vaccine effectiveness in the long term and against newly emerging variants of the coronavirus. In this context, it is imperative to assess the risks of disease transmission due to various common activities to enable individuals to make informed decisions about engagement. Air travel is one such activity.

Although highly symptomatic COVID-19 carriers are unlikely to be on commercial aircraft due to current airline travel policies, asymptomatic and presymptomatic carriers continue to travel, and mildly symptomatic carriers who are able to board undetected do so with some regularity. According to contact tracing organizations [1], this often occurs when the subject becomes infected while away on travel, and travels while potentially symptomatic in order to return home. Nonetheless, reports of COVID-19 transmission onboard aircraft are rare, with no confirmed cases for domestic travel within the U.S. at the time of writing, despite 1,600 cases of potentially symptomatic travelers that have been investigated by the Center for Disease Control and Prevention [1]. Since contact tracing is difficult when travel is involved due to the decentralized structure of the current efforts, the present study was performed in order to complement the epidemiological data.

The expiratory activity selected for study was a cough because it is a higher-magnitude perturbation than either breathing or talking in terms of both the number and the volume of expiratory particles, as well as in terms of momentum, which was expected to translate to a greater challenge to particle removal by the aircraft. Sneezing, which is a higher-magnitude perturbation than coughing, was not selected for study because it is not a symptom of COVID-19 [2]. The Computational Fluid Dynamics (CFD) model that had been developed in the aftermath of SARS-CoV-1 with funding from the Federal Aviation Administration [3,4,5,6] was updated to reflect improvements in the understanding of expiratory emissions and in the availability of computational resources. Analysis was performed to predict material transport during and after a cough discharge. Material inhaled by other passengers was used to quantify passenger exposure – or, viewed alternately, to characterize the efficiency of the aircraft system at protecting passengers from exposure to infectious material.

### Airflow Design

While the air distribution system on modern aircraft provides far better protection against airborne pathogens than the vast majority of other common environments, its design has been motivated by the comfort of passengers and the flight crew and the cooling requirements of the on-board electronics systems.

Under most conditions, air supplied to the aircraft cabin is a mixture of outside air with air removed from the cabin, which is filtered before it is recirculated. Recirculation is used both to improve fuel efficiency and to increase humidity during flight, as humidity is a factor in comfort. A less common condition that may occur during ground operations is that no outside air is available, so all air supplied to the cabin is the filtered recirculated air. Either scenario provides high-purity air through the use of high-efficiency particulate air (HEPA) filters, which remove 99.97% of particles at 0.3 μm [7], a filtration level that is sufficient to remove viruses contained in droplet nuclei of respiratory emissions. The outside air is assumed to be free of pathogens, and its pressure, temperature, and humidity are adjusted to comfortable levels on the way to the cabin by either the Environmental Control System or the ground supply sources.

Air is introduced into the passenger cabin through air distribution nozzles, which are typically located above the seats to maximize thermal comfort for passengers and the crew. On the 737 interior used in the current study, air distribution nozzles are located outboard of the Passenger Service Unit and direct the airflow inboard toward the center of the cabin, Fig. 1. The supply airstream remains attached to the PSU and the stow bin through the Coandă effect [8].

**Fig. 1.**
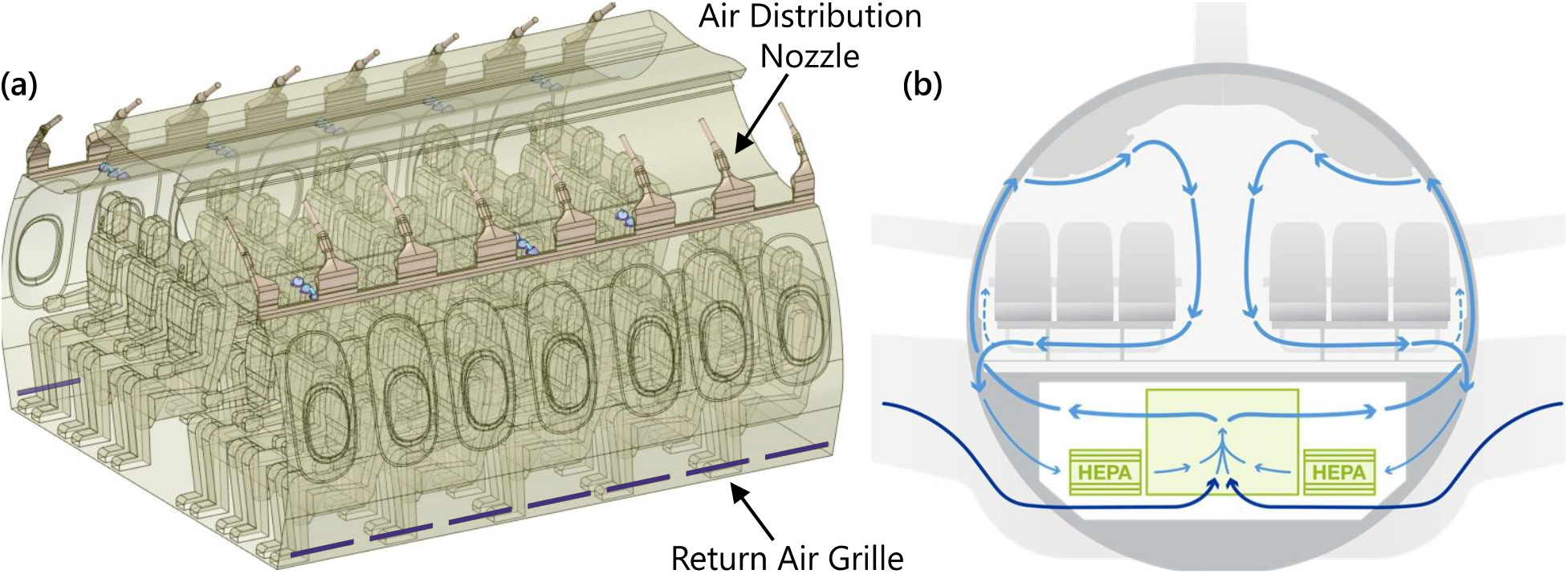
Airflow design in narrow-body aircraft. **(a)** Airflow delivery features in the 737 Boeing Sky Interior cabin used in simulations. **(b)** Idealized airflow pattern in the passenger cabin.

On reaching the aisle, the airstream combines with the opposing airstream and in the process redirects toward the floor, producing higher velocities in the aisle compared to the seated sections, which compartmentalizes the left and right sections into cells. At the floor, the stream splits and continues outboard to either side of the cabin and toward the return air grilles, with low pressure behind the grilles driving the motion. Air which does not exit through the grilles rises by buoyancy due to heat sources from the passengers and the In-Flight Entertainment Systems and becomes entrained in the high-velocity jet at the air distribution nozzle. Then the process repeats again.

The resulting airflow pattern consists of two counter-rotating cells formed around the rows of passenger seating. High seat backs support the cohesion of the cell structure and limit fore-aft movement. This compartmentalized design was originally developed during the era when in-flight smoking had been the norm as a means to reduce exposure to second-hand smoke of the passengers and the flight crew. Since cigarette smoke has particles in a similar size range as expiratory emissions, this design was also expected to limit the spread of potentially infectious particles in the era of COVID-19.

### Simulation Overview

Simulations were performed in a 5-row, 30-seat section of the 737 Boeing Sky Interior cabin with periodic front and back interfaces, Fig. 1(a). Forward-facing passengers occupied all seats. Personal Air Outlets were left off due to the large number of permutations of available PAO positions and the absence of a recommendation for PAO use in the COVID-19 context. A breathing zone was defined around the face of each passenger to track particle exposure, shown in Supplementary Fig. S2. The size of each breathing zone was 1 ft on each side or 0.8 ft^3^ due to the volume occupied by each passenger’s face.

Supplementary Table S1 summarizes the simulated cases, which varied the seat assignment of the coughing index passenger, the airflow rate, and the randomly generated initial condition. One airflow rate was studied with initial conditions from three different time points to reflect the right-to-left shifts in the counter-rotating cell structure that occur on a periodic basis, as shown in Fig. 2. Separate initial steady-state solutions were required for each airflow rate and are indicated in the case table.

**Fig. 2.**
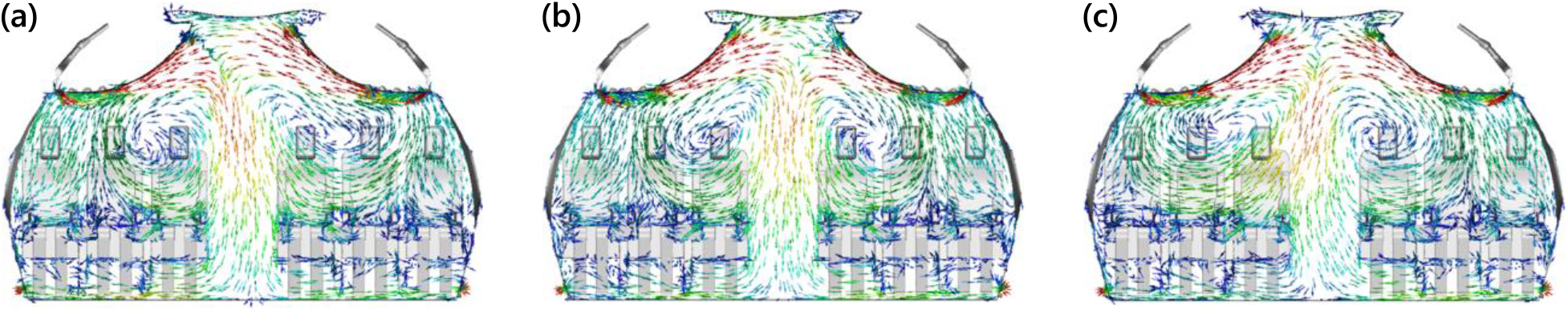
Velocity vectors of initial conditions at 100% flow rate with the highest velocities shown in red and the lowest ones in blue. Time offset and initial condition index as follows: **(a)** 0 s, Initial Condition 1, **(b)** 90 s, Initial Condition 2, **(c)** 120 s, Initial Condition 3.

## Results

### Particle Removal Dynamics

Overall dynamics of particles expelled by the cough were tracked by following the decay in particle count over the course of each simulation, Fig. 3. The initial features on the decay curves corresponded to deposition onto surfaces such as the seat back in front of the index subject. Approximately 50% of the nonvolatile material was deposited onto various surfaces, with the rest removed through the return air grilles located near the floor on the interior walls.

**Fig. 3.**
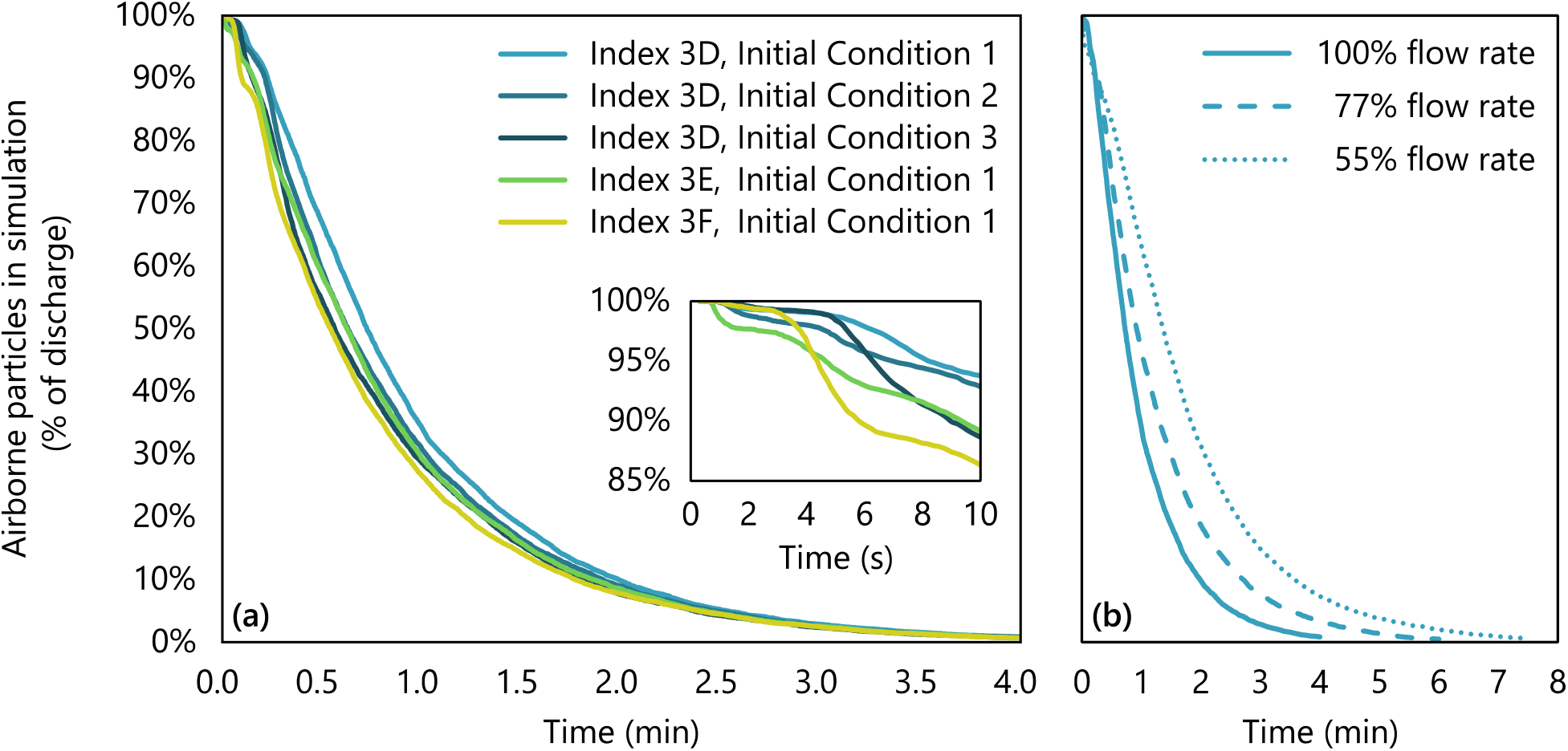
Decay of expiratory particles over time after cough discharge. **(a)** Airflow at 100% with different index seats and initial conditions. **(b)** Airflow at 55-100% with index passenger in aisle seat 3D.

Differences due to random fluctuations of the flow field were captured effectively by differences in starting conditions. These random differences affected the rate of particle removal from the cabin over the first 1-2 minutes, Fig. 3(a), but their effect decreased over time, and the final time for particle removal was independent of the initial condition. For an index passenger in seat 3D with airflow at 100%, initial conditions resulted in only a 10-second difference in the time to remove 95% of the particles from the cabin.

Supply airflow rate, on the other hand, had a large effect on the rate of particle removal, Fig. 3(b). At 100% flow rate, 95% of the particles were removed in 2.4 to 2.5 minutes (80% in 1.3 to 1.4 min, 99% in 3.7 to 3.8 min). However, at 55% flow rate, it took 4.6 minutes to reach 95% removal of particles (80% in 2.6 min, 99% in 6.9 minutes).

Particle dynamics in the breathing zones of susceptible passengers were tracked by following the particles by mass over time, Fig. 4. All particles were dehydrated by the time they reached the nearby breathing zones, so their masses are expressed as a percentage of the original nonvolatile content. Particle decay in a given breathing zone by mass was faster than the overall rate of particle removal by particle count. For example, for an index passenger in middle seat 3E, susceptible passenger in aisle seat 3D, and 100% airflow, 95% of the cumulative nonvolatile mass was removed in 1.4 minutes (80% in 0.7 min, 99% in 2.3 min). A seat chart is provided in Fig. 5.

**Fig. 4.**
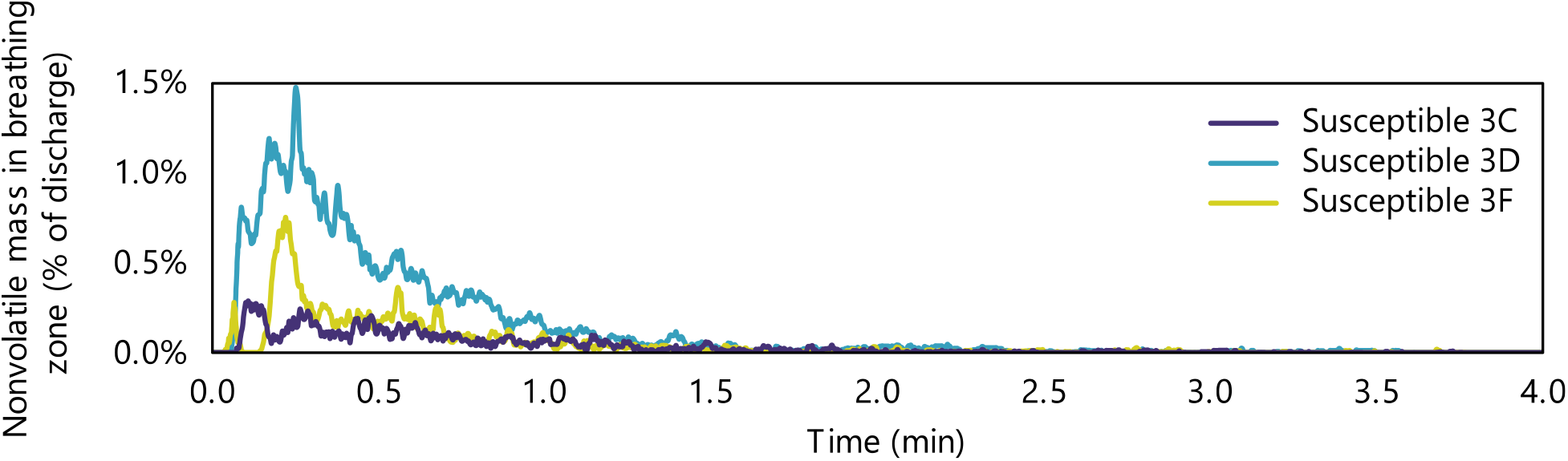
Amount of expiratory material present in the breathing zones of susceptible passengers over time after cough discharge from an index passenger in middle seat 3E.

**Fig. 5.**
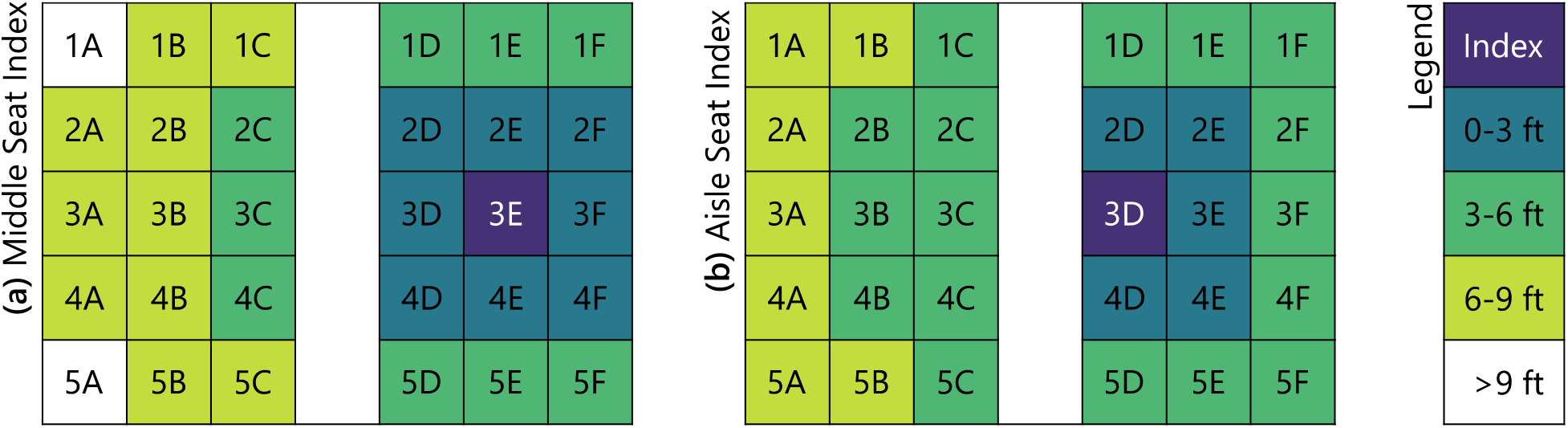
Seat chart for the 5-row cabin section used in the model, colored by distance of susceptible passengers from an index passenger seated in either **(a)** the middle or **(b)** the aisle seat.

### Particle Exposure

Inhaled mass in each susceptible passenger’s breathing zone was integrated over the course of each simulation, with cumulative exposures presented in Fig. 6. Although proximity to the index subject was only a weak predictor of exposure, Supplementary Fig. S10, in aggregate, passengers seated closer to the index were exposed to more index expiratory material than those farther away. Exposure was highest for passengers seated in the index subject’s row, and lowest for passengers seated two rows away. Exposure was higher on the right side of the cabin, where the index passenger’s seat was located, than on the opposite side. This spatial distribution pattern was consistent with the counter-rotating cell pattern of airflow depicted schematically in Fig. 1(b) and Fig. 2.

**Fig. 6.**
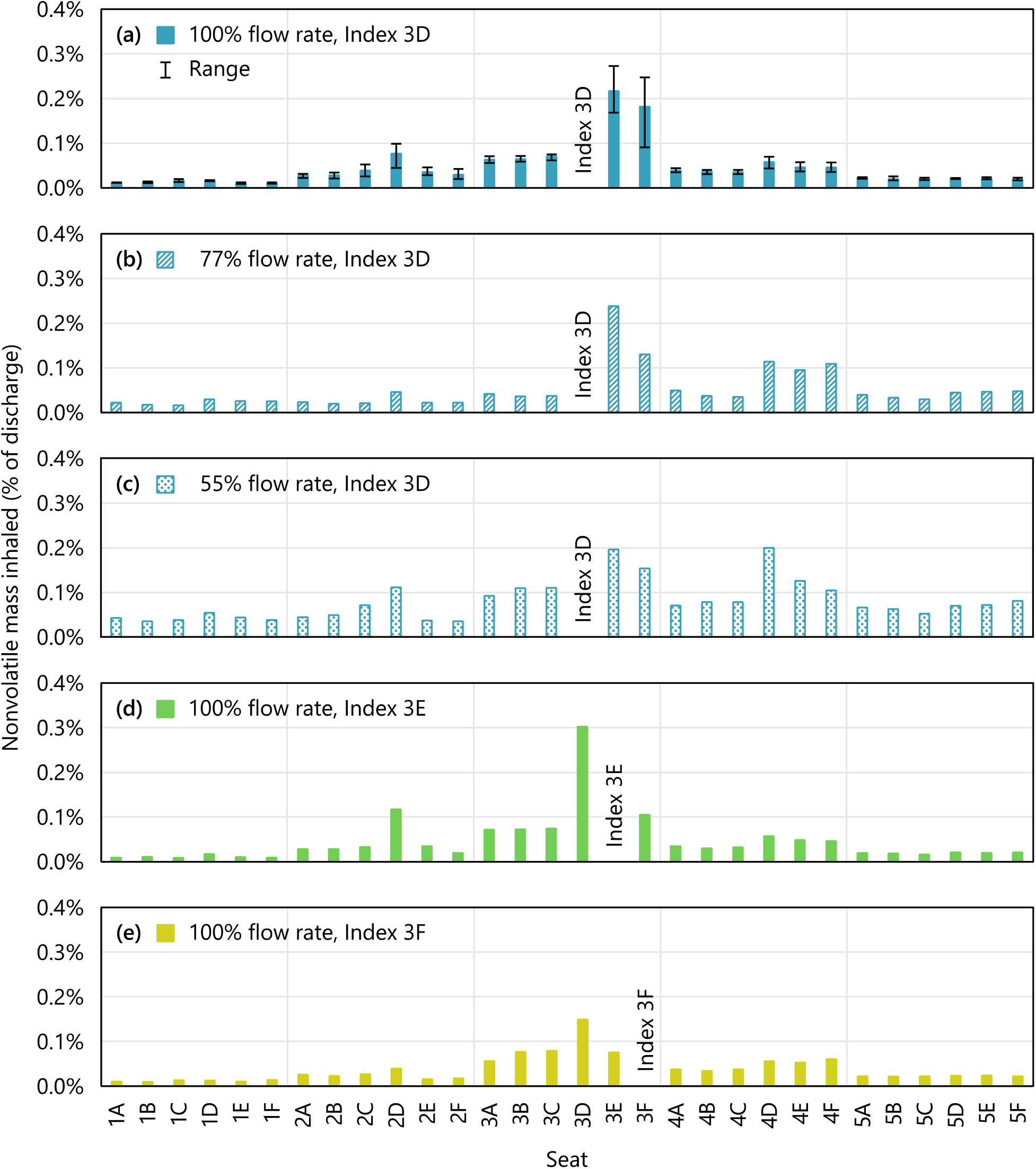
Mass of expiratory material inhaled by susceptible subjects in different seats for **(a-c)** different airflow conditions and **(a, d-e)** different index seat assignments at 100% airflow. Range bars in (a) represent values over three different initial conditions.

Exposure of susceptible passengers in the index subject’s row was lowest when the index subject was in the window seat. Exposure of passengers in the window seat was lower than exposure of passengers in the aisle seat. However, all exposures were a small fraction of the cough discharge: the maximum mass inhaled by a susceptible passenger was 0.3%, Supplementary Fig. S6. This occurred for the susceptible passenger in seat 3D when the index passenger was in seat 3E, with dynamics shown in Fig. 4 and Supplementary Fig. S5. Differences due to random fluctuations of the flow field had a substantial effect on exposure for some of the seats, with the maximum coefficient of variation of 45%, which occurred for the susceptible passenger in seat 3F. The minimum coefficient of variation was 5.4% and the median was 15%.

Reducing airflow from 100% to 55% resulted in a wider spread of particles throughout the cabin, observed as an increase of mass inhaled by most susceptible passengers regardless of their proximity to the index subject, Supplementary Fig. S9. However, the maximum exposure across all susceptible passengers did not increase. For some passengers, exposure decreased as airflow was reduced, which is attributed to random differences in initial conditions.

## Discussion

Transport of particles expelled by a cough was studied to characterize the effectiveness of airplane ventilation and airflow patterns in protecting passengers from exposure to an infected index subject. This case was studied both as a perturbation to quantify the efficiency of the aircraft system in removing particles from the passenger cabin, and as a scenario that currently commands public interest.

The amount of expiratory material inhaled by susceptible passengers was quantified in terms of nonvolatile mass and expressed as a percentage of the nonvolatile mass discharged by the index subject. This material transport approach is not specific to any disease, and was used because the input data required for a deterministic disease transmission model of SARS-CoV-2 are not currently available. Specifically, the distribution of SARS-CoV-2 across particles of different sizes is not known, but is likely non-uniform: for example, influenza RNA is shed predominantly in smaller particles [9,10,11] even though larger particles account for the bulk of the expelled volume. Further, smaller particles may be more infectious, as they deposit deeper in the respiratory tract when inhaled, and fewer pathogens are needed with other diseases to produce infection in the lower vs. upper respiratory tract [12,13,14]. These factors are important because the size of particles affects their aerodynamic behavior, so the absence of inputs specific to SARS-CoV-2 limits the extent of persuasive analysis.

Viewed alternatively, the current material transport approach is equivalent to making the simplifying assumptions that the pathogen of interest is distributed uniformly across particles of different sizes, and that the infectious dose is independent of particle size. Given those assumptions, the exposure of susceptible subjects in terms of a percentage of nonvolatile mass expelled would be equivalent to the percentage of viruses expelled, which could then be related to virus shedding rates and infectious doses to extend the present work to a disease transmission model. That approach was not used here because of the uncertainty that these simplifying assumptions are valid.

Nonetheless, a key finding of the present study was that exposure to respiratory particles expelled by the index passenger appeared low even for the passenger’s nearest neighbors, with a maximum exposure of 0.3% of the nonvolatile mass expelled. The amount inhaled by the susceptible passengers depended primarily on their proximity to the index subject and on airflow rate, with random fluctuations of the flow field also having a significant role. Humidity did not have a substantial effect within the low-humidity range that is typical of cruise altitudes, Supplementary Figs. S7 and S8.

Three airflow cases were studied that cover the range of airflow rates that would be expected over the course of a journey, with 77% being the average. The lowest flow rate studied, 55%, may occur on the ground during the boarding and deplaning segments, when both the engines and the auxiliary power unit (APU) are off, and no ground air supply is available. While this scenario is less common, it provided a lower bound airflow rate for calculating exposures of susceptible passengers to an infected index person. Decreasing the airflow rate increased the dispersion of expiratory particles throughout the cabin, but did not increase the maximum exposure to expiratory material. While exposures of passengers seated away from the index subject increased, exposures of the index passenger’s neighbors remained the same within the variability caused by random flow field fluctuations. For passengers seated away from the index subject, exposure remained on par or lower than exposure of passengers seated in the same triplet of seats as the index.

### Experimental Comparison

The present results are consistent with the results of recent experimental testing [15,16,17,18,19,20,21] that was funded by the United States Transport Command (TRANSCOM) Air Mobility Command (AMC) division of the U.S. Military, and was planned and carried out by a large team that included some of the authors of the present paper. In that work, tracer aerosols were discharged from various seat locations throughout Boeing 767 and 777 passenger cabins, both on the ground and in flight, and their distribution over time was measured from 40+ nearby seats for each discharge. A subset of 75 test discharges whose conditions most closely matched those in the computational work was used for comparisons. Data were binned by distance to the index subject as shown in Fig. 5 to compare the model to the experimental results in both the near and the intermediate field, and to account for differences in seating configurations.

Particle exposures were within the same order of magnitude, Fig. 7. While the highest exposure values were observed experimentally, those values were outliers, which were far less likely to be captured in the computational work due to the much smaller sample size: only one computational case was used for comparisons because it was the only case that matched the experimental airflow conditions. On the other hand, the median values were surprisingly similar given the differences in methods that were used to generate the two datasets, Table 1. A partial explanation is that the metric that was used to describe results, exposure as a percentage of discharge, appears sufficiently robust to account for the differences in discharge volumes. A second explanation is that some variables that were expected to affect exposure had no appreciable effect, such as the discharge velocity, which was varied experimentally in a subset of tests, Supplementary Fig. S11.

**Fig. 7.**
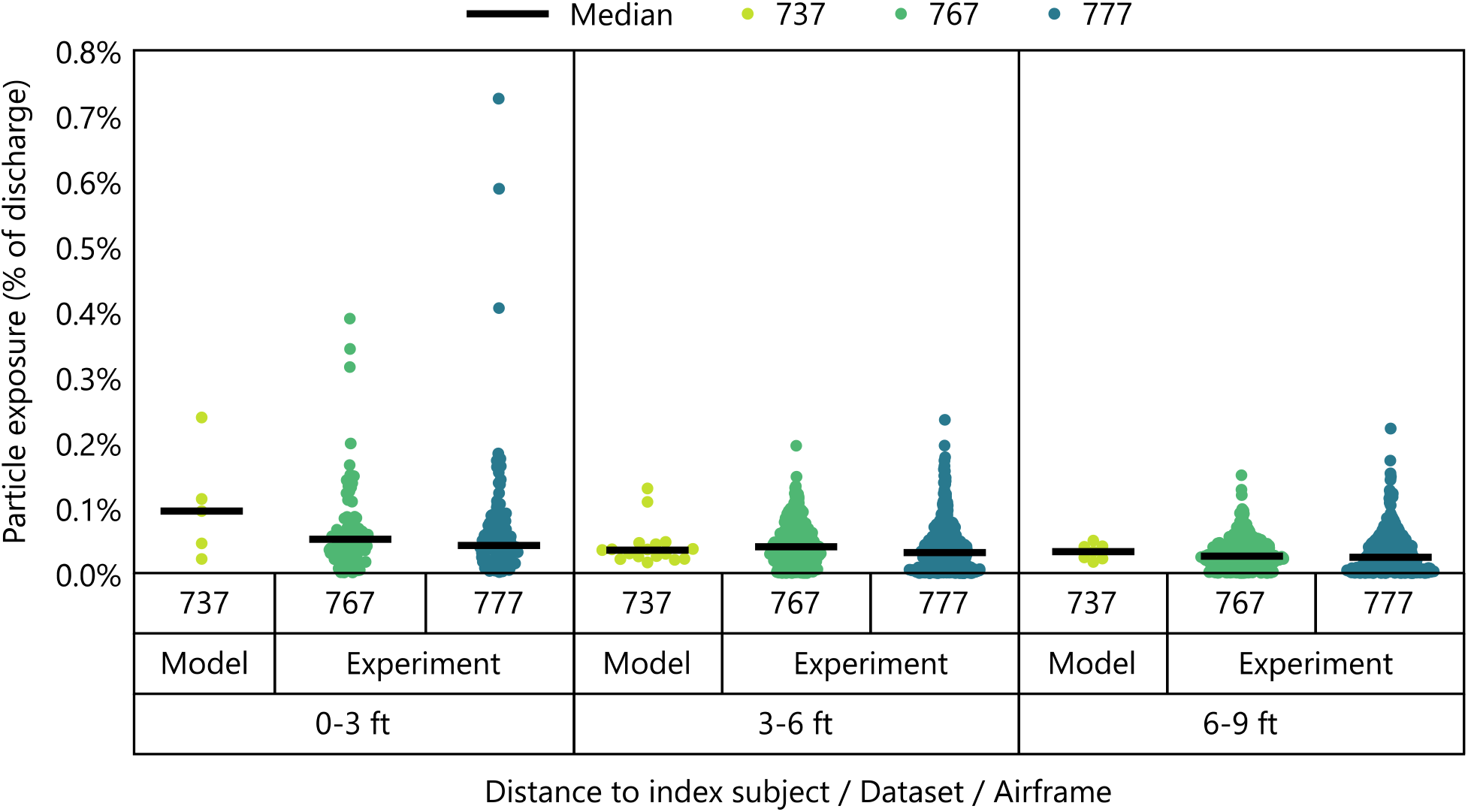
Particle exposure in the computational model using the 737 airframe and in experimental testing using 767 and 777 airframes. Jittered datapoints represent exposures of individual susceptible passengers within bins by distance to the index subject.

**Table 1.** Comparison of methods and results used to characterize particle exposure in the computational model and in experimental testing performed by the TRANSCOM/AMC team.

|  | Property | Model | Experiment |
| --- | --- | --- | --- |
| Aircraft | Airframe | Boeing 737-800 NG or Boeing 737-8 MAX | Boeing 777-200 and Boeing 767-300 |
|  | Airframe type | Narrow-body | Wide-body |
|  | Airflow rate | 77% | Similar to 77% |
| Expiratory Material | Size distribution | Polydispersed, based on voluntary coughs of healthy volunteers | Narrowly dispersed around 2.7 $\mu\text{m}$ before evaporation and 1 $\mu\text{m}$ after [15,22] |
|  | Nonvolatile fraction | 10% | 5% |
|  | Interactions with surfaces | 100% adhesion on contact | Some adhesion on contact |
| Passengers | Discharge volume | 0.054 $\mu\text{L}$ before and 0.0054 $\mu\text{L}$ after evaporation | 1.9 $\mu\text{L}$ before and 0.094 $\mu\text{L}$ after evaporation |
|  | Discharge duration | 0.4 seconds | 1 minute |
|  | Peak discharge velocity | 9.2 m/s | 1.4 m/s |
|  | Heat output per subject | 70 W | 15 W |
| Results | Exposure, Maximum | 0.24% | 0.73% |
|  | Exposure, Median within 3 ft of index | 0.10% | 0.04% |
|  | Exposure, Median 3-6 ft from index | 0.03% | 0.03% |
|  | Exposure, Median 6-9 ft from index | 0.03% | 0.03% |

The main difference in results was the effect of distance. While median exposure values were similar between the model and the experimental testing for seats more than 3 ft from the index location, median exposure was higher in the model for susceptible seats that were within 3 ft. At the time of writing, this difference in trends cannot be attributed to any specific methodological factor since the effects of several methodological variables on the spatial distribution of particles have not been characterized in prior work. First, the airflow structures of the single-aisle aircraft used in the model and the twin-aisle aircraft used experimentally are inherently different, but how that difference affects particle transport is unknown. Second, the size of particles affects their aerodynamic behavior, and different size distributions were used in the model and experimentally to represent either coughing or breathing. However, most studies of the relationship between travel range and particle size are performed in the absence of airflow, so the effect on transport in high airflow conditions that are characteristic of aircraft but rare otherwise is also unknown.

Using data reported here, passenger exposure cannot be resolved by particle size because of the manner in which simulations had been set up. This constitutes a limitation of the present work, not only because it has bearing on the experimental comparison, but also because particles of different sizes are likely to have different pathogenicities [12–14] – and the present data does not allow to calculate exposure levels for particles in different size fractions, or by particle count or surface area. Therefore, it is notable that exposure levels from the model are on par with those from experimental testing because particles that had been used experimentally would be small enough to reach the lower respiratory tract [12] where the consequences of particle deposition are likely to be most severe. That supports the relevance of the present approach of quantifying exposure by the percentage of mass.

### Limitations

Cases studied in the present work were intended to be representative of the typical cabin environment. However, the effects of most of the factors that are expected to affect material transport have not been studied, either here or elsewhere. The following uncharacterized effects are expected:

- The cabin configuration affects the airflow pattern, which varies somewhat throughout the cabin. The cabin configuration includes selections made by the airline operator, such as the seating arrangement for First and Business class and the class divider, galley separators, and lavatory locations. It also includes temporary modifications made by the passengers, such as the use and position of Personal Air Outlets, the position of tray tables and stow bins, the reclined or upright seat positions, the presence of under-seat luggage, and the level of passenger loading. In addition, seat geometry and pitch may affect the particle content in the breathing zones of susceptible subjects.
- The on-ground cabin environment defined by the operator and the airport, including temperature and humidity, is expected to affect particle dynamics and may affect the airflow pattern. During boarding and deplaning, the aircraft engines must be turned off, and the APU may be required to be turned off by airport regulations, such that the operator is responsible for providing air conditioning via ground-based sources. When no efficient ground-based conditioning is available, temperature and humidity may deviate significantly from the values studied. (However, recirculated air continues to be provided at the 55% level included in this study.)
- Further, during boarding and deplaning, the entry door is open and the passengers and flight crew move around the cabin, which significantly alters the airflow pattern. Subject movements also occur during meal service and to use the lavatories. In addition to airflow, subject movements change the spatial relationships between index and susceptible subjects, as do variations in the passengers’ seated postures.
- The use of masks and other face coverings capture a fraction of the expired material and redirect the flow of the remainder.

These factors combine to create infinite permutations. However, an exhaustive case exploration is not required to generalize the results of the current study. Data from the present model shows that some exposure onboard aircraft can occur, so vulnerable populations are advised to remain vigilant; but, the level of exposure appears to be low under the most representative circumstances, consistent with both the experimental work by TRANSCOM/AMC [15–21], and the absence of confirmed transmissions due to air travel within the U.S. [1].

## Methods

### Particle Generation

The cough aerosol was modeled using the particle count, size distribution, and material volume obtained by Zayas *et al*. [23], which is consistent in shape with the dataset from Morawska *et al*. [24]. While a wide range of distributions and concentrations exists in the current literature, the Zayas *et al*. dataset was selected because it was developed using a fast-acquisition and high-resolution laser diffraction system that enabled the team to identify very small particles at a high particle density. In contrast, many literature reports have a minimum bound of 0.37 μm which, based on the Zayas *et al*. dataset, would have the effect of undercounting 82% of particles, even if the lower bound were due to simple truncation.

Of the two distributions reported by Zayas *et al*., the dataset without the outlier (superemitter) was used. Superemission was not considered in this study; however, due to the need for computational simplification described in the following section, the choice of a superemitter would not have affected results. Absolute particle counts were obtained from the metric reported by Zayas *et al*. of particles per cm^3^ using the instrument analysis volume of 7.85 cm^3^. This brought the count to approximately 106M for a reported material volume of 0.0544 μL. (In contrast, some current literature reports include particle counts in the tens of thousands). No correction was made to account for the distance from the mouth of the test subject to the instrument sensor, which was 17 cm. The resulting particle size distribution is shown in Supplementary Fig. S3.

Cough dynamics were modeled using the methodology of Gupta *et al*. [3]. The arbitrarily selected inputs of cough peak flow rate of 3.67 L/s, cough expired volume of 0.7 L, and peak velocity time of 76.2 ms were used to generate the profile of airflow expelled by the cough, which was truncated at 0.4 seconds (or 10% of the cough peak flow rate) and is shown in Supplementary Fig. S4. A mouth opening area of 4 cm^2^ was used as the value for an average male, and a single cough angle of 27.5° was used as the midpoint of the 95% confidence interval values provided by Gupta *et al*.

### Particle Dynamics

Computational Fluid Dynamics modeling was conducted using Ansys Fluent 19.2 software. A 737 cabin section with dimensions in Supplementary Fig. S1 was meshed using 13.3M polyhedral cells. Thermal boundary conditions, listed in Supplementary Table S2, were selected to maintain the cabin temperature at the passenger-preferred 75°F while applying the airflow rates of interest. Composition of the continuous phase (the mixture of air and water vapor) was solved using the Eulerian scheme, which accounted for interactions between expiratory particles and the surrounding medium, including heat transfer and evaporation. In this scheme, expiratory particles were modeled as water with a volatile fraction of 90% [25]. However, their vapor pressure was adjusted by a factor of 0.28 to account for the effect of pulmonary surfactants [26]. Using this approach, particles evaporated gradually over about 1 second until reaching 10% of their initial mass.

Steady-state solutions were used as the initial conditions for the subsequent transient simulations. A time step of 0.05 seconds was used after the initial impulsive event which was calculated using shorter time steps. Transport equations in the continuous phase were solved using the realizable *k-ε* turbulence model with enhanced wall treatment. Transport equations of expiratory particles were solved using the Lagrangian discrete phase model. To reduce computational complexity, the volume of each cough was scaled down by a factor of 157 from a total of 106M particles to 676,208, and was scaled back at the end of the simulations for post-processing. Particles were tracked in parcels, or statistical representations of individual droplets, of an average of 19.57 particles each, for an average of 34,555 parcels per simulation. On contact with surfaces, particles deposited with complete efficiency to account for the effect of pulmonary surfactants [27,28]. The overall CFD analysis of each case required thousands of cores in parallel for several days.

### Particle Inhalation

Particles were assumed to have evaporated to their nonvolatile fractions by the time they reached the nearby breathing zones. This is a conservative assumption: if inaccurate, reported values would overestimate actual exposures. As the first post-processing step, the nonvolatile volume of particles in each breathing zone was determined by taking the total mass of particles and converting it to volume using the density of water.

For each passenger, the inhalation portion of a sinusoidal tidal breathing curve was calculated as follows:

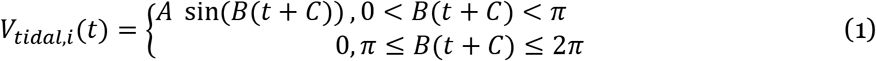

Using equations provided by Gupta *et al*. [4], *A* was taken as 0.616 L/s and *B* as 2.23 s^−1^ by assuming an average U.S. male with a weight of 89.7 kg and a height of 1.75 m [29]. This was a conservative choice, since the minute volume calculated for an average U.S. male is about 25% higher than that for an average U.S. female. The phase shift *C* was varied for each passenger to obtain the maximum nonvolatile volume that could have been inhaled. No provision was made to re-exhale [12] a fraction of the inhaled material. An example inhalation curve is shown in Supplementary Fig. S5.

The well-mixed model was applied to each breathing zone as a simplifying assumption that is not expected to bias results. The nonvolatile volume of particles in the breathing zone at a given time, *V_bz,p_*(*t*), was divided by the total 22.7 L volume of the breathing zone, *V_bz,tot_*, to get the nonvolatile particle fraction in the breathing zone, which was multiplied by the time-dependent tidal inhalation volume *V_tidal,i_*(*t*), and then integrated over the time of the simulation *t_s_* as follows:

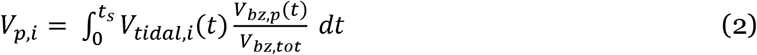

Results of example calculations are shown in Supplementary Fig. S5.

Data analysis and plotting were performed in Excel 2013, MATLAB R2019b, and JMP14.

### Experimental Comparison

Experimental measurements by the TRANSCOM/AMC team had been carried out under several sets of experimental conditions [15–21]. The following subset of conditions was used to compare results to the present model:

- Masks and Personal Air Outlets were off to match conditions used in the computational work.
- The insert that was used to accelerate the discharge velocity to a cough-representative value was off. The number of tests that used this modification was small and its use did not qualitatively affect results, Supplementary Fig. S11, so this condition was excluded to reduce the number of variables across the different datasets. (Supplementary Fig. S11 was produced by comparing tests where the velocity modification insert was the only variable.)
- Economy seating sections in the aft and mid-aft of the aircraft were pooled for the analysis. Seating configurations in the premium sections included features that were different from the model, including partitions between passengers which were up during testing, and seats that were positioned at an angle rather than facing forward. Data from premium sections was not included in the comparison to more closely match conditions in the computational work.
- In-flight testing at altitude and ground-based testing with the aircraft door closed were pooled for the analysis. Ground-based testing with the aircraft door open was not included in the comparison to match conditions in the computational work. The number of releases affected by the exclusion was small and this choice did not qualitatively affect results, Supplementary Fig. S12.
- Door-closed test 777-2 which included data generated during a partial equipment failure was excluded from analysis to reduce the number of variables.
- Door-open test 777-36 whose duration was uncertain due to experimental error was excluded as well.

All comparisons were against the simulation with 77% airflow rate because that rate most closely matched the experimental conditions.

Data were compared in terms of particle exposure expressed as a percentage of the amount discharged. For the model, analysis was based on the nonvolatile mass of expiratory material that was inhaled by susceptible subjects because metrics other than mass had not been collected during simulations. For experimental measurements, it was based on particle counts that were scaled to account for the difference between the instrument sampling volume and the minute volume of susceptible subjects used in the model. Since the particles used experimentally were dispersed narrowly by their size, particle counts were nearly equivalent to nonvolatile masses when expressed as a percentage of amount discharged, which enabled the comparisons despite the different metrics.

Comparisons were performed as a function of distance between representations of index and susceptible passengers. For the model, distances were calculated using mouth and nose xy coordinates of the manikins. For experimental testing, they were based on an idealized placement of the nebulizer outlet and the particle collector ports. Error in calculated distances due to non-ideal placement of testing instrumentation as well as any discrepancy between estimated and actual seat coordinates is estimated as less than 1 ft. Calculated distances were binned into increments of 3 ft and described using medians. The bin size was chosen for convenience as the choice had no qualitative effect on results, Supplementary Fig. S13. Medians were chosen to describe the data because they were less sensitive to outliers than means, Supplementary Fig. S14, and because sample sizes were small relative to the number of possible cabin configurations.

## Supporting information

Data used to generate all data figures

## Data Availability

All inputs and outputs reported in this paper are provided either in the main text or as Supplementary Information, which includes both graphical representations and Excel files of tabular data. Researchers are encouraged to contact the corresponding author for clarifications, advice on data utilization, and collaborations.

## Acknowledgements

The authors thank Chao-Hsin Lin, Matthew J. Schwab, and Robert J. Atmur for many productive discussions on the Environmental Control System, CFD modeling and the content of this paper. The authors thank the High Performance Computing staff at Boeing who prioritized access to computing resources for this activity and provided around-the-clock server support. The authors acknowledge Ansys for providing licenses for their Fluent software to support Confident Travel Initiative studies at Boeing.

## Author Contributions

ACD, ADC, TW, SPJ, LLW, and NAO designed the experiments and set the methodologies.

ACD, ADC, TW, and NAO provided parameterization prior to TW and ADC performing computational fluid dynamics calculations.

ACD, MZ, ADC, TW, and NAO performed data analysis and validated the data content.

ACD, MZ, and NAO curated the data.

ACD, ADC, TW, and NAO wrote sections of the original draft and edited the document.

MZ wrote and edited the original draft and all subsequent versions of the paper.

ADC and JJC provided resources including access to off-the-shelf software, software licenses, configuration parameters, labor, and computational resources.

JJC provided supervision, administration, and acquisition of funding.

NAO as corresponding author ensured that original data upon which the submission is based is preserved and retrievable for reanalysis, approved data presentation as representative of the original data, and has foreseen and minimized obstacles to the sharing of data described in the work.

