## Supplemental info and figures for "Computational Fluid Dynamics Modeling of Cough Transport in an Aircraft Cabin"

#### File Contents

Supplementary Methods ..................... S3

Supplementary Tables ..................... S4

**Table S1.** Case conditions varied in simulations. ..................... S4

**Table S2.** Thermal boundary conditions used in simulations. ..................... S4

Supplementary Figures ..................... S5

**Fig. S1.** Dimensions of the 737 Boeing Sky Interior section used in simulations. ..................... S5

**Fig. S2.** Breathing zone definition in front of the face of each susceptible passenger used in simulations. ..................... S6

**Fig. S3.** Particle size distribution of respiratory material expelled by the simulated cough. ..................... S7

**Fig. S4.** Rate of airflow expelled by the simulated cough. ..................... S7

**Fig. S5.** Sample calculations of particle inhalation. ..................... S8

**Fig. S6.** Spatial distribution of expiratory material inhaled by susceptible passengers. ..................... S9

**Fig. S7.** Decay of expiratory particles over time, including supplementary cases at 10 and 20% RH. ..................... S10

**Fig. S8.** Spatial distribution of expiratory material inhaled by susceptible passengers for supplementary cases at 10 and 20% RH. ..................... S10

**Fig. S9.** Particle exposure of susceptible subjects using different airflow rates and index seat assignments. ..................... S11

**Fig. S10.** Relationship between particle exposure of susceptible subjects and their distance from the index subject. .......... S12

**Fig. S11.** Particle exposure of susceptible subjects in computational simulations and in experimental testing that varied particle discharge velocities to model different expiratory activities of the index subject. .......... S13

**Fig. S12.** Particle exposure of susceptible subjects in experimental testing performed either in flight, at an airport hangar with the aircraft door closed, or at an airport terminal with the aircraft door open. .......... S14

**Fig. S13.** Particle exposure of susceptible subjects binned by distance to the index subject using increments of either 2, 3, or 4 ft. .......... S15

**Fig. S14.** Particle exposure of susceptible subjects summarized using both medians and means. .......... S16

Supplementary References .......... S17

### Additional Supplementary Materials

Tabular Data

**Data for Fig. S3.** Particle size distribution of expiratory material.

**Data for Fig. S4.** Airflow rate of simulated cough.

**Data for Fig. 4.** Mass of expiratory material in breathing zones of sample susceptible passengers.

**Data for Fig. S5.** Sample calculations of particle inhalation.

**Data for Figs. 3 and S7.** Number of particles in simulation over time.

**Data for Figs. 6, S6, and S8.** Particle exposure of susceptible passengers.

**Data for Fig. S9.** Particle exposure of susceptible passengers binned by distance to the index subject.

**Data for Figs. 7 and S14.** Particle exposure of susceptible passengers in simulation compared to experimental measurements.

### Supplementary Methods

The effect of ambient humidity on particle transport was evaluated in combination with the effect of breathing by susceptible passengers. Humidity in the continuous phase was introduced into the cabin by increasing the relative humidity of the supply nozzle and through passenger respiration. The supply nozzle was set to either 11% or 25% relative humidity (RH) to reach an average RH of 10% or 20% at cabin temperature, Table S1. Passengers exhaled air at 31°C and 100% RH.

Respiration was modeled using the methodology of Gupta *et al.* [1]. Breathing was performed through the nose with a nostril area of 0.71 $cm^2$, a front breathing angle of 60°, and a side breathing angle of 69°. For each passenger, the tidal breathing curve was calculated as:

$$V_{tidal}(t) = A\ \sin(B(t + C)) \tag{S1}$$

Random values of $A$ and $C$ were generated for all susceptible passengers, with $A$ ranging from 0.34 to 0.42 L/s. As a consequence of the random phase shift, passengers were out of phase in their breathing activities. The same period with $B$=0.45π $s^{-1}$ was used for all passengers.

### Supplementary Tables

**Table S1.** Case conditions varied in simulations.

| Flow Rate (%) | Flow Rate (ACFM) | Initial Condition Index | Offset from Initial Condition 1 | Index Passenger Seat | Relative Humidity (%) |
|---|---|---|---|---|---|
| 100 | 588 | 1 | 0 s | 3D | 0 |
| 100 | 588 | 2 | 90 s | 3D | 0 |
| 100 | 588 | 3 | 120 s | 3D | 0 |
| 77 | 453 | 4 | N/A | 3D | 0 |
| 55 | 323 | 5 | N/A | 3D | 0 |
| 100 | 588 | 1 | 0 s | 3E | 0 |
| 100 | 588 | 1 | 0 s | 3F | 0 |
| 100 | 588 | 6 | N/A | 3D | 10 |
| 100 | 588 | 7 | N/A | 3D | 20 |

**Table S2.** Thermal boundary conditions used in simulations.

| Type | Value |
|---|---|
| Heat output per passenger | 70 W |
| Heat flux of stowage bins, ceiling, and floor | Adiabatic wall |
| Sidewall temperature | 55-65°F |
| Supply nozzle temperature | 62-67°F |
| Average cabin temperature | 75-77°F |

### Supplementary Figures

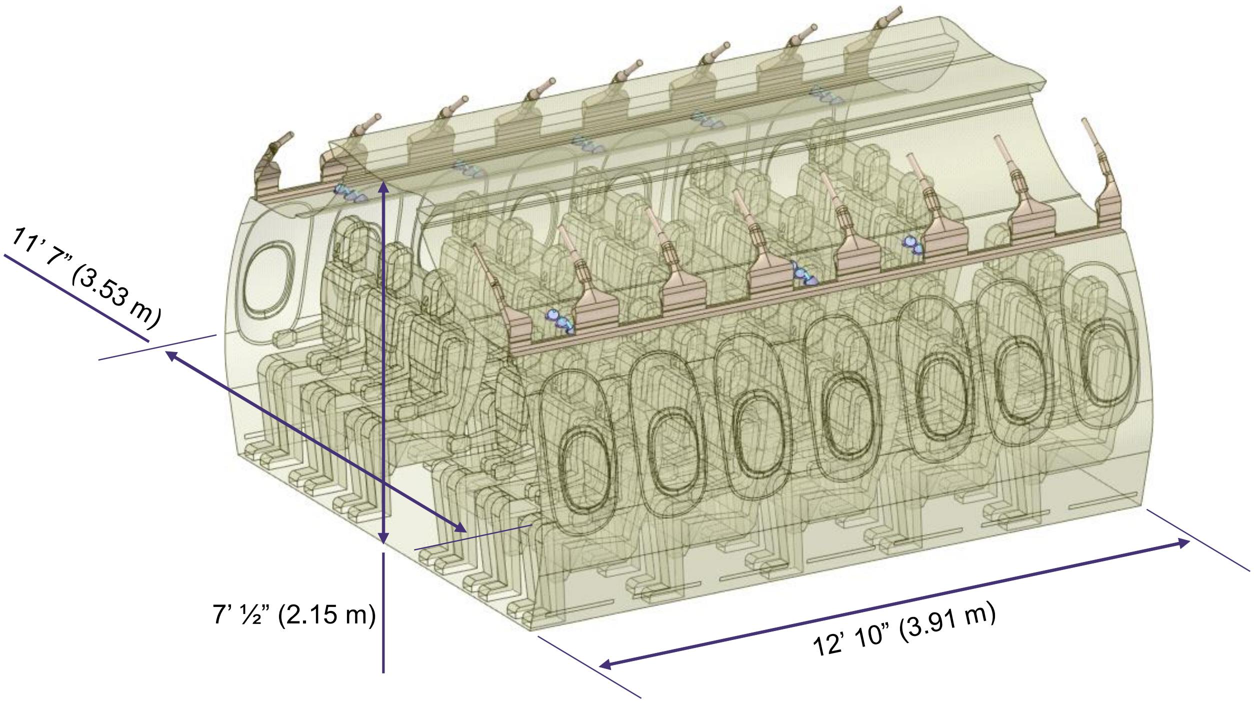


**Fig. S1.** Dimensions of the 737 Boeing Sky Interior section used in simulations.

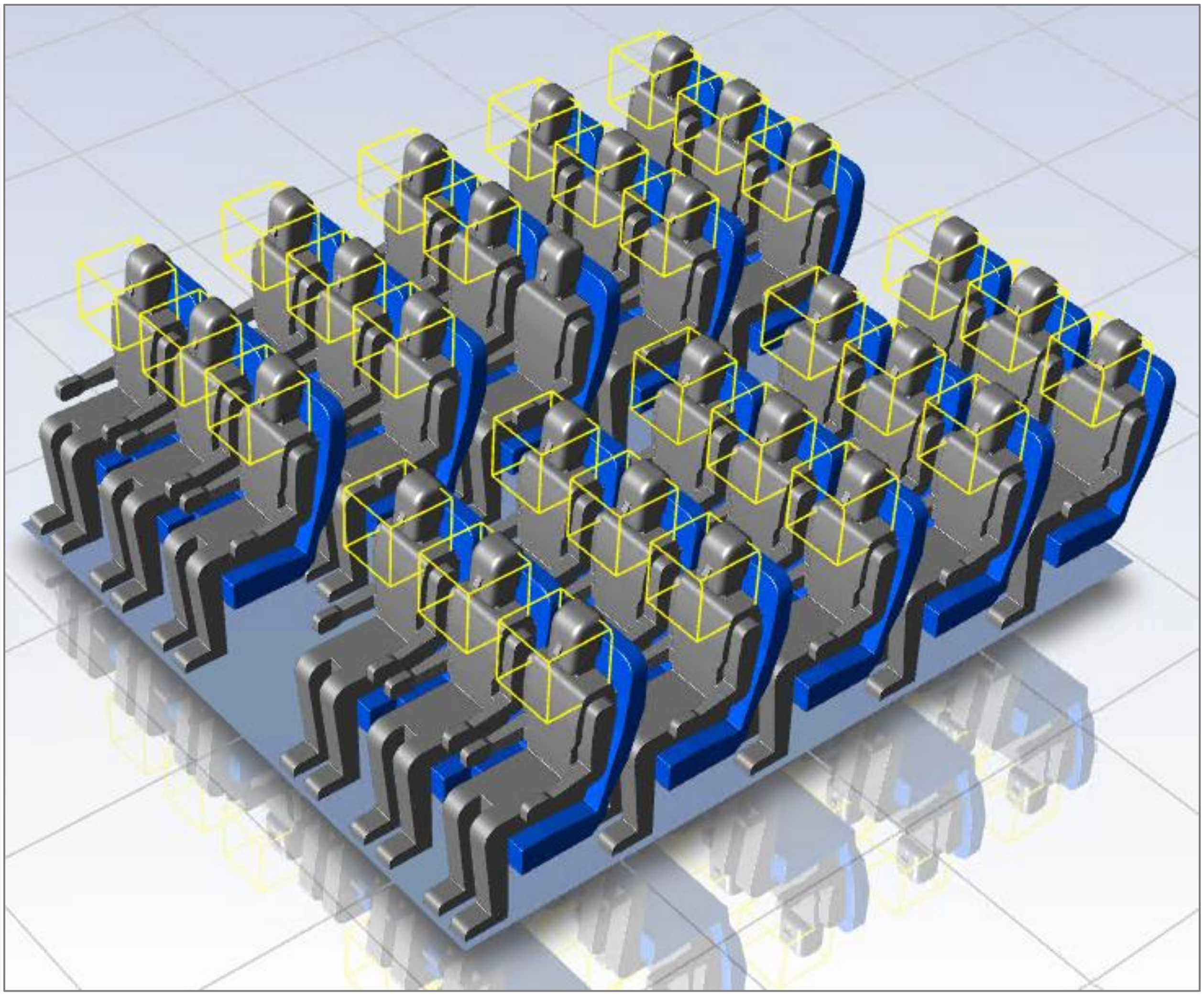

**Fig. S2.** Breathing zone definition in front of the face of each susceptible passenger used in simulations, shown with the example of index passenger in aisle seat 3D.

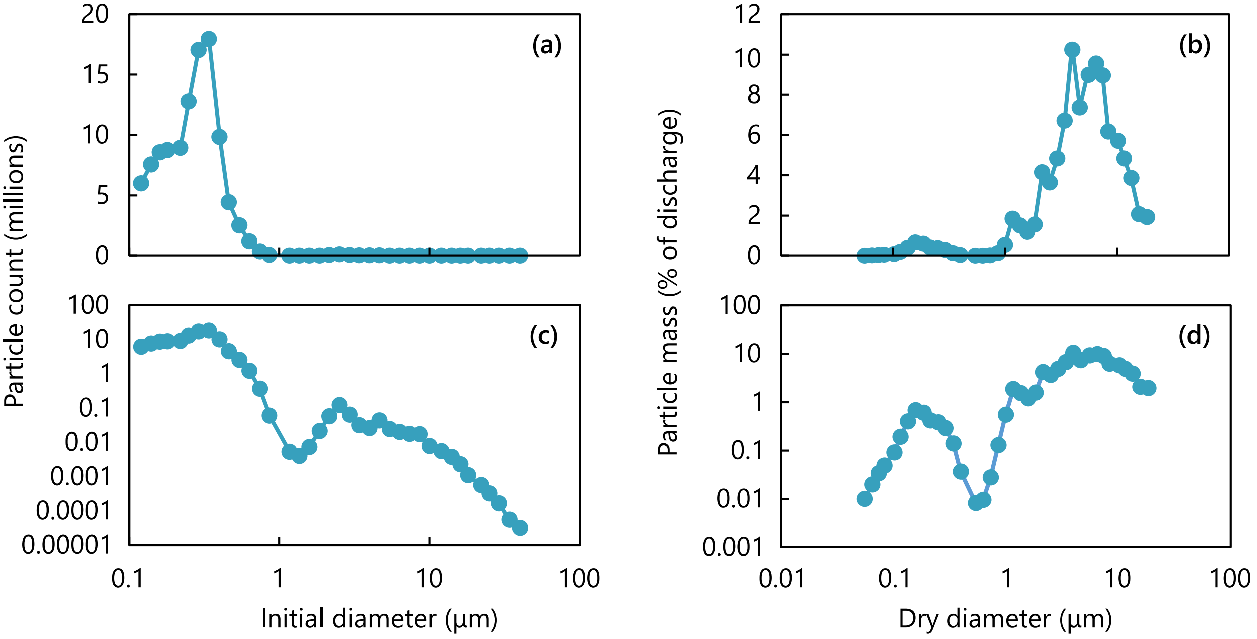


**Fig. S3.** Particle size distribution of respiratory material expelled by the simulated cough, shown on **(a,b)** lin-log and **(c,d)** log-log scales, and either in terms of **(a,c)** particle count for each diameter at the time of discharge, or **(b,d)** total particle mass for each diameter after drying out to 10% of the initial mass. Adapted from Zayas *et al.* [2].

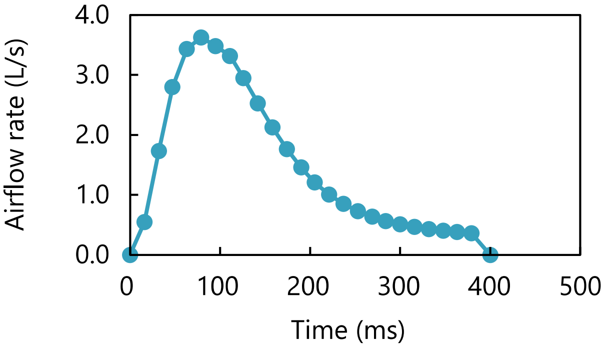


**Fig. S4.** Rate of airflow expelled by the simulated cough. Adapted from Gupta *et al.* [3].

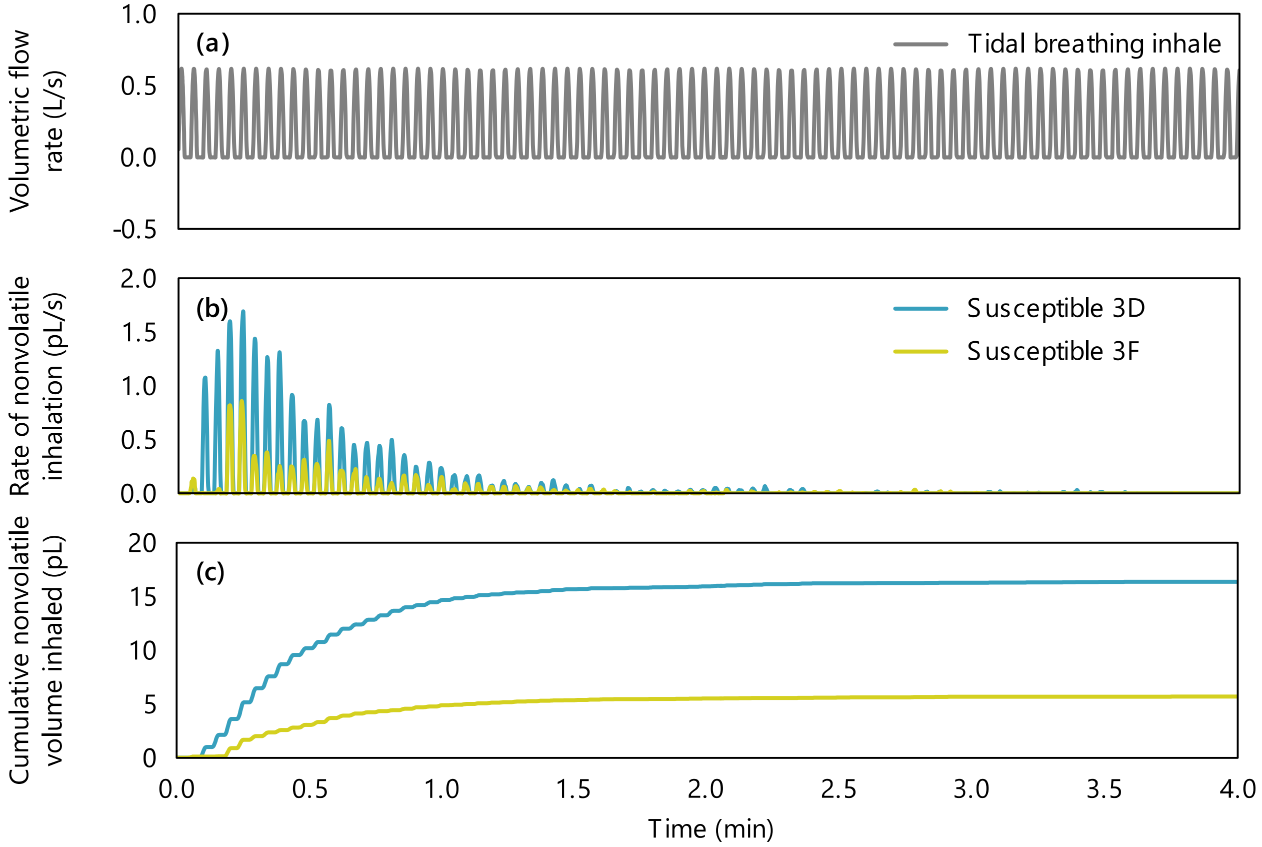


**Fig. S5.** Sample calculations of particle inhalation for index passenger in seat 3E. **(a)** Inhalation portion of the tidal breathing curve before adjusting phase shift. **(b)** Nonvolatile volumetric inhalation rate and **(c)** cumulative nonvolatile volume inhaled versus time for susceptible passengers in aisle seat 3D and window seat 3F.

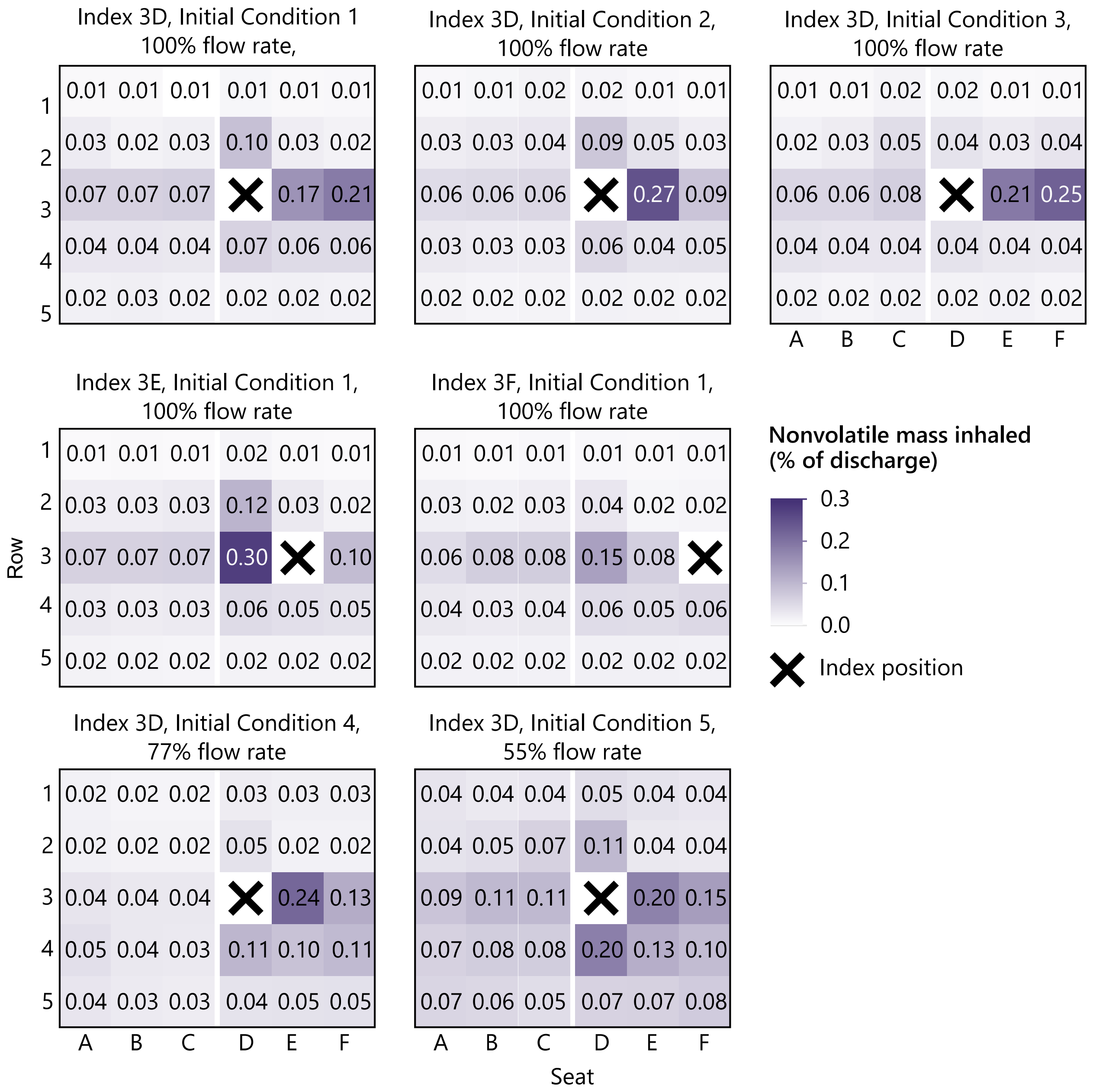


**Fig. S6.** Spatial distribution of expiratory material inhaled by susceptible passengers.

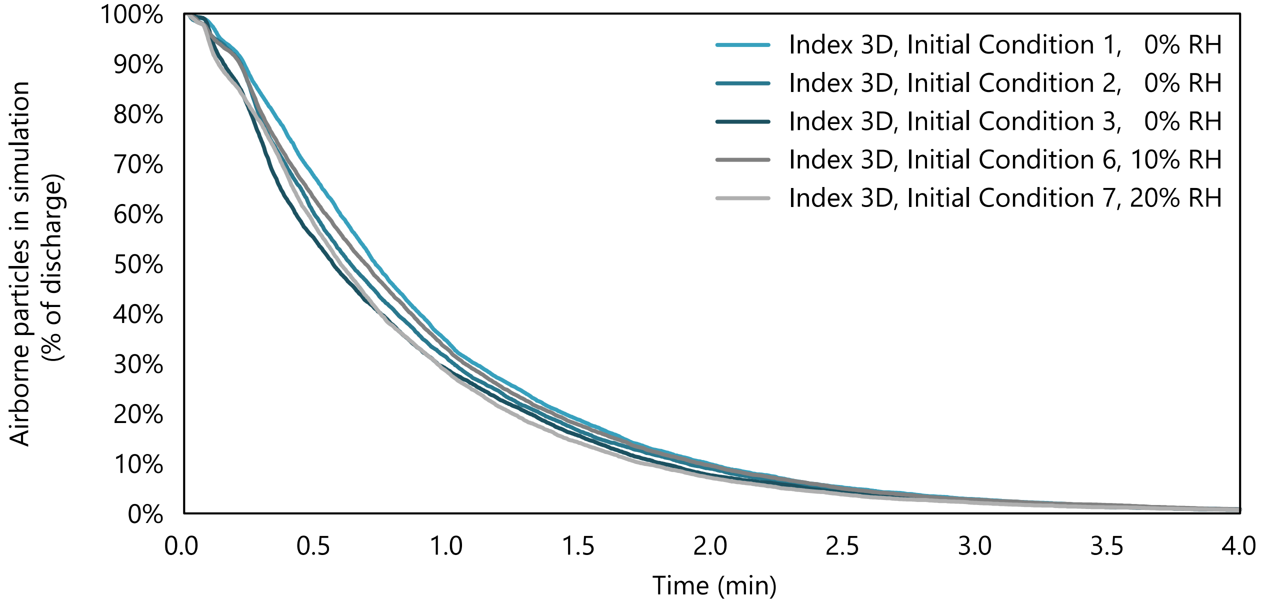


**Fig. S7.** Decay of expiratory particles over time after initial discharge at 100% airflow with different index seats and initial conditions, including supplementary cases at 10 and 20% RH.

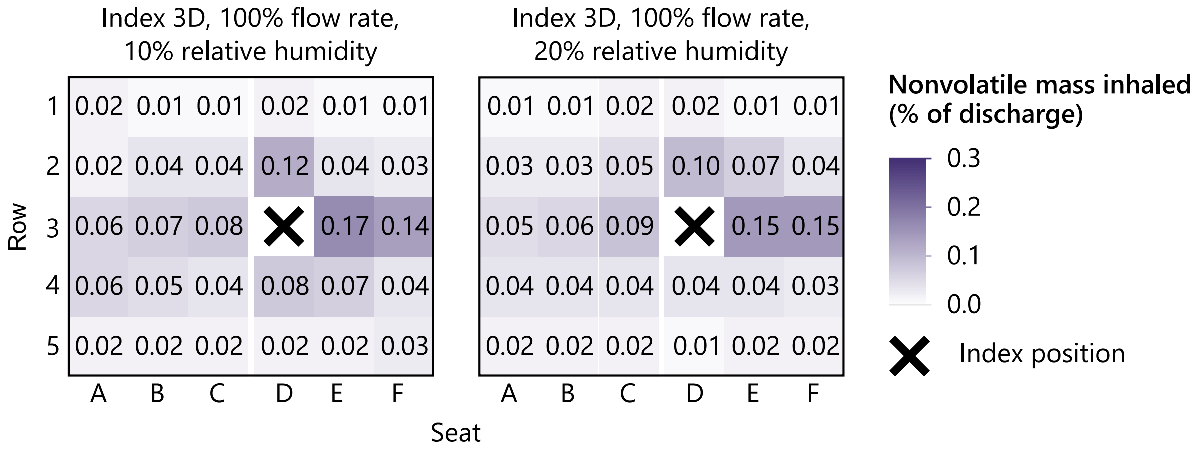


**Fig. S8.** Spatial distribution of expiratory material inhaled by susceptible passengers for supplementary cases at 10 and 20% RH.

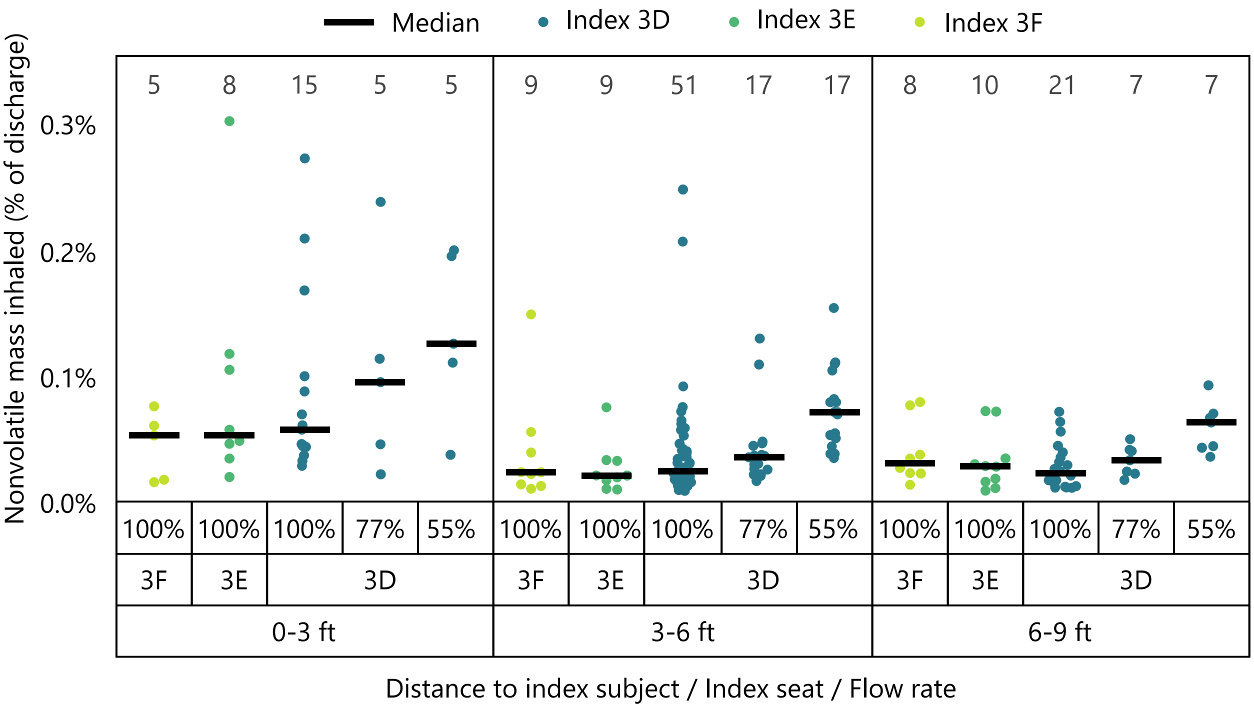


**Fig. S9.** Particle exposure of susceptible subjects using different airflow rates and index seat assignments. Jittered datapoints represent individual subject exposures with the number of datapoints indicated above each bin.

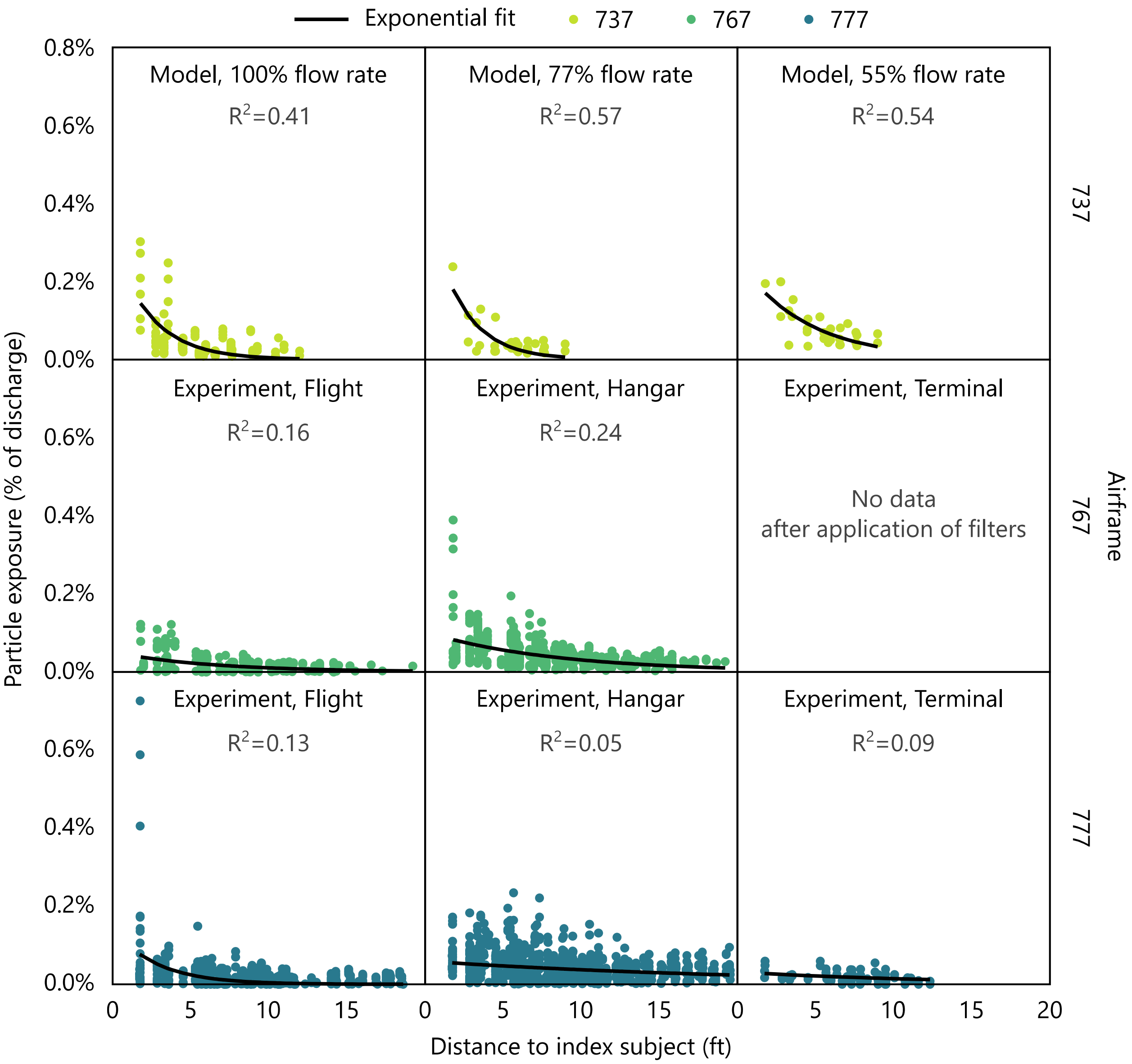


**Fig. S10.** Relationship between particle exposure of susceptible subjects and their distance from the index subject.

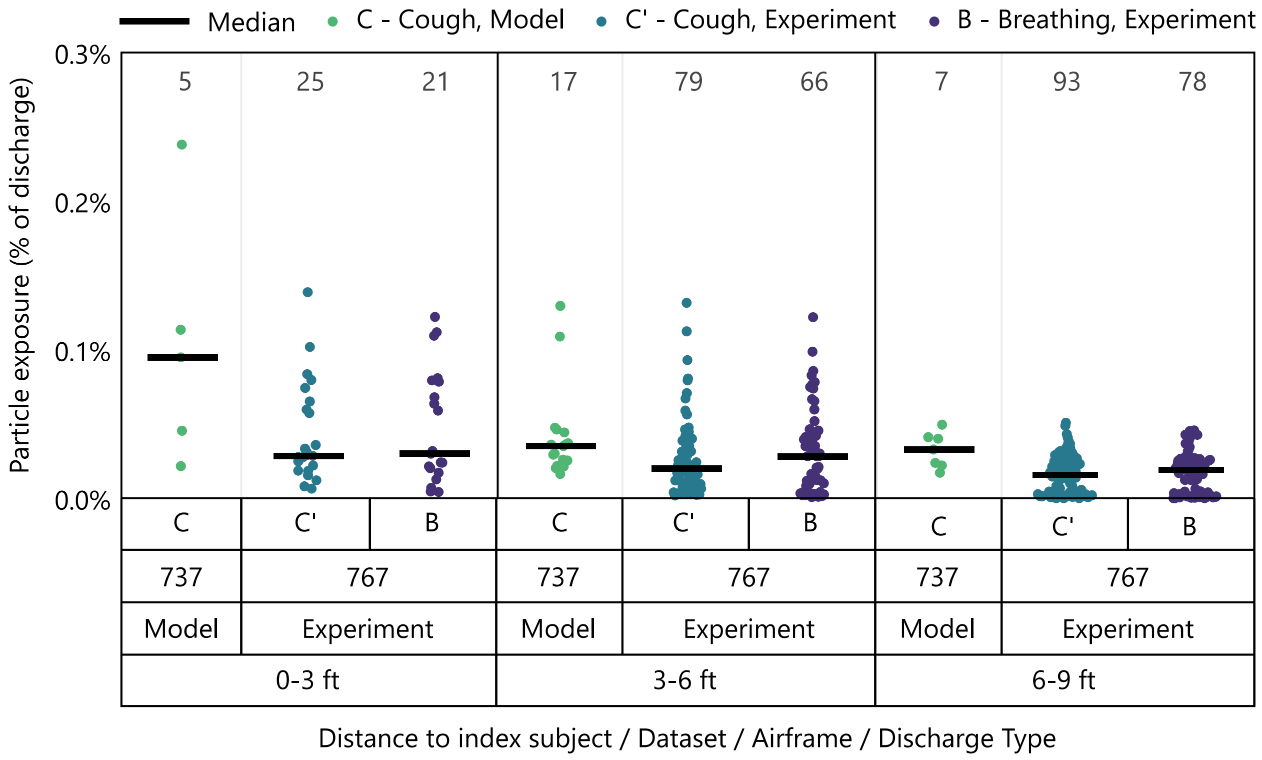


**Fig. S11.** Particle exposure of susceptible subjects in computational simulations and in experimental testing that varied particle discharge velocities to model different expiratory activities of the index subject. Jittered datapoints represent exposures of individual subjects with the number of datapoints indicated above each bin.

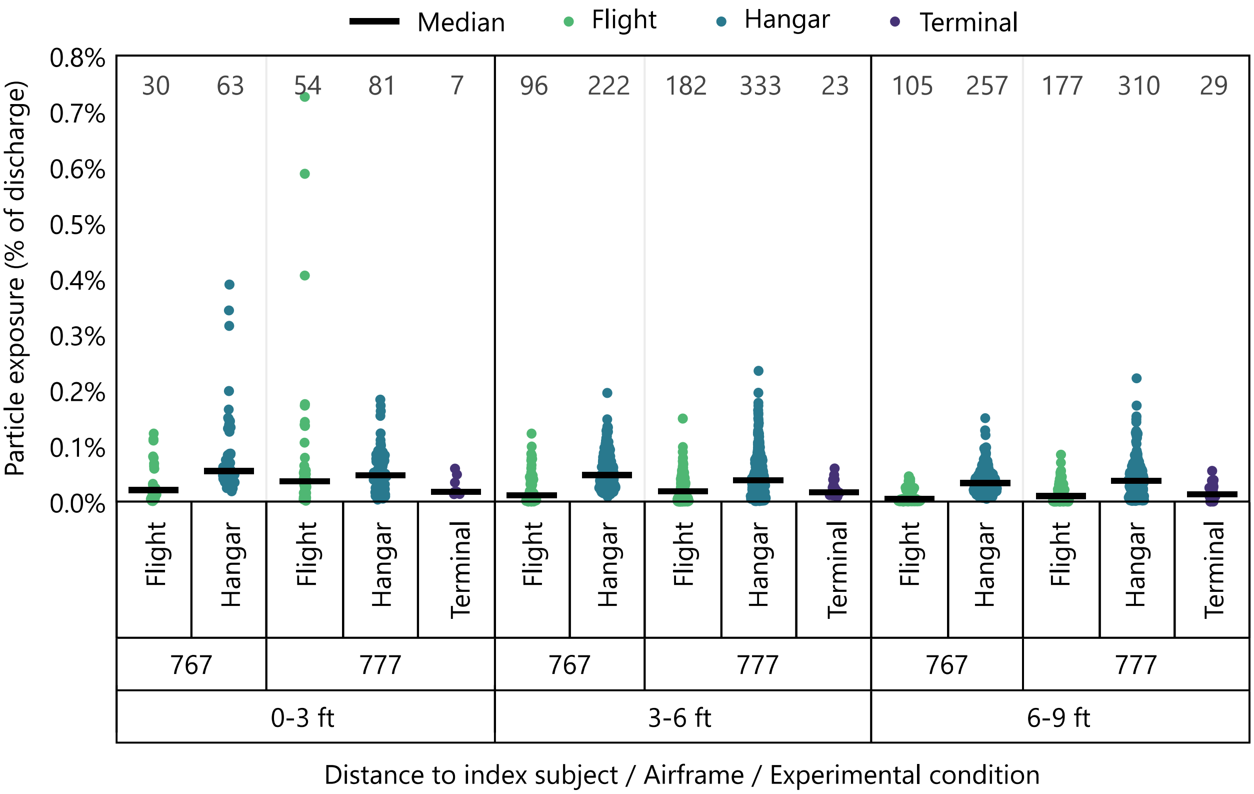


**Fig. S12.** Particle exposure of susceptible subjects in experimental testing performed either in flight, at an airport hangar with the aircraft door closed, or at an airport terminal with the aircraft door open. Jittered datapoints represent exposures of individual subjects with the number of datapoints indicated above each bin.

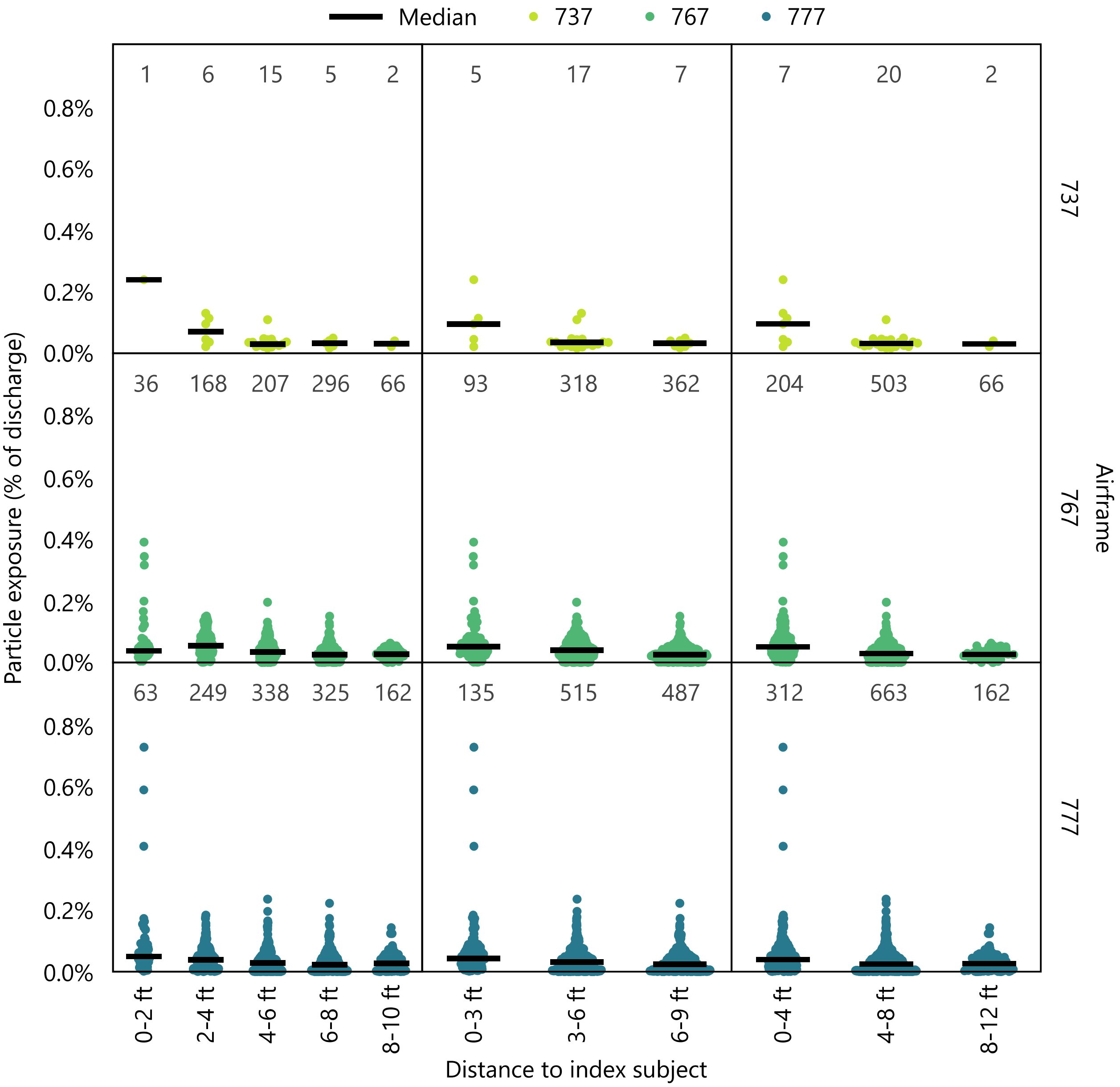


**Fig. S13.** Particle exposure of susceptible subjects binned by distance to the index subject using increments of either 2, 3, or 4 ft. Jittered datapoints represent individual subject exposures with the number of datapoints indicated above each bin.

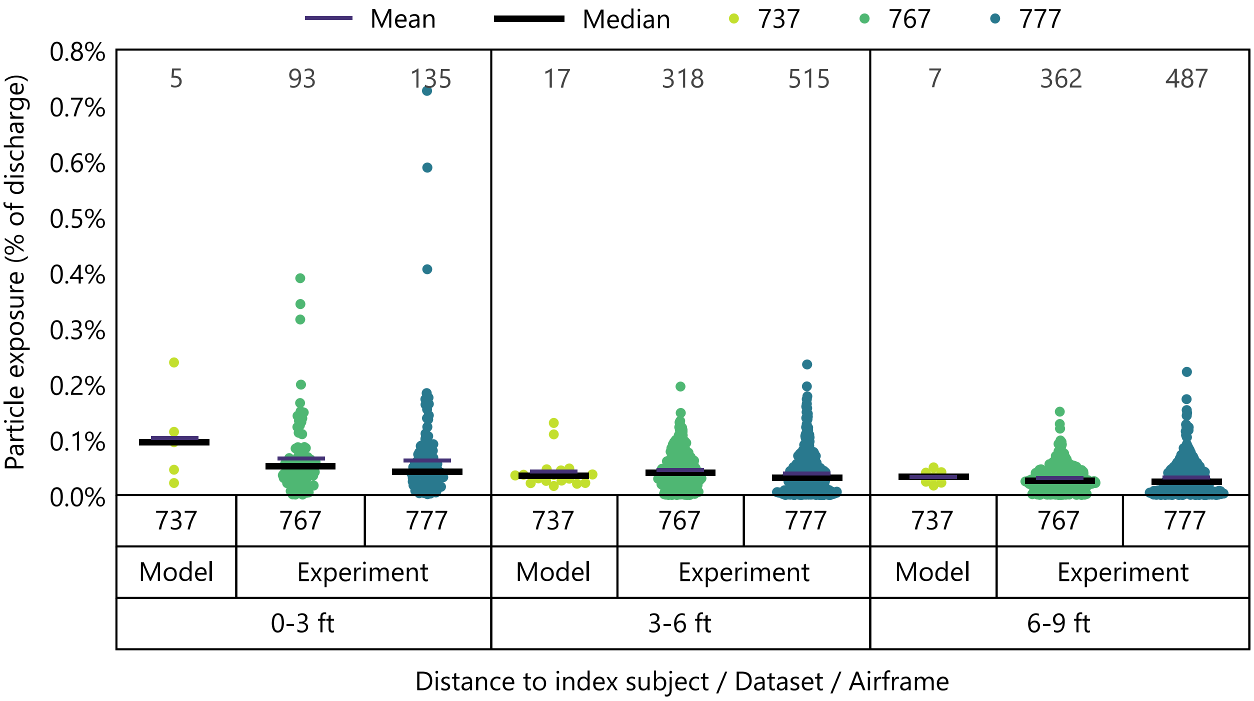


**Fig. S14.** Particle exposure of susceptible subjects summarized using both medians and means. Jittered datapoints represent individual subject exposures with the number of datapoints indicated above each bin.
